# SINAPS: Prediction of microbial traits from marker gene sequences

**DOI:** 10.1101/124156

**Authors:** Robert C. Edgar

## Abstract

Microbial communities are often studied by sequencing marker genes such as 16S ribosomal RNA. Marker gene sequences can be used to assess diversity and taxonomy, but do not directly measure functions arising from other genes in the community metagenome. Such functions can be predicted by algorithms that associate marker genes with experimentally determined traits in well-studied species. Typically, such methods use ancestral state reconstruction. Here I describe SINAPS, a new algorithm that predicts traits for marker gene sequences using a fast, simple word-counting algorithm that does not require alignments or trees. A measure of prediction confidence is obtained by bootstrapping. I tested SINAPS predictions from 16S V4 query sequences for traits including energy metabolism, Gram-positive staining, presence of a flagellum, V4 primer mismatches, and 16S copy number. Accuracy was >90% except for copy number, where a large majority of predictions were within +/−2 of the true value.

## Introduction

Next-generation sequencing has revolutionized the study of microbial communities in environments ranging from the human body (Cho & Blaser 2012; Pflughoeft & Versalovic 2012) to oceans (Moran 2015) and soils (Hartmann et al. 2014). There are two main approaches in such studies: *marker gene metagenomics*, in which a single gene such as 16S ribosomal RNA is amplified, and *shotgun metagenomics* in which DNA from a sample containing microbes is cleaved into random fragments. Shotgun metagenomics could, in principle, reconstruct the complete genome of all species in a sample, but in practice is limited to assembling at most a few of the most abundant species. Lower-abundance genomes are represented by short fragments which often cannot be reliably assigned a taxonomy or function. Amplifying marker genes enables consistent detection of low-abundance species, providing a more complete characterization of the taxonomic content of the community without giving a direct indication of functions represented by other genes in the community metagenome. This motivates the development of methods for predicting functional traits from marker gene sequences. The central challenge confronting such methods is that only a few thousand complete microbial genomes are currently available, necessitating extrapolation of functional annotations to the large majority of species known only by their marker gene sequences. A natural strategy for extrapolation is *ancestral state reconstruction*, which examines a tree for the marker gene in which leaf nodes with known traits are annotated. If a trait is seen to be conserved within the known leaves of a subtree, then other species in that subtree are inferred to have the same trait. One method that uses ancestral state reconstruction is PICRUSt (Langille et al. 2013) which infers genes and pathways classified by the COGS (Tatusov et al. 1997) and KEGG (Ogata et al. 1999) databases, respectively. A similar approach has been used to predict 16S copy number (Kembel et al. 2012). Ancestral state reconstruction is intuitively appealing, but is complicated to implement and can be computationally expensive. Also, a given implementation may introduce undesirable limitations; for example, PICRUSt uses “closed-reference” OTU clustering which discards reads having < 97% identity with the subset of Greengenes that is used as a reference, potentially losing many or even most species in a community with many novel taxa.

Taxonomy prediction methods can be interpreted as extrapolation algorithms where the trait is membership of a given named clade. From this perspective, the taxonomy assignment methods (McDonald et al. 2012; Yilmaz et al. 2014) of Greengenes (DeSantis et al. 2006) and SILVA (Pruesse et al. 2007) respectively use ancestral state reconstruction, while the RDP Classifier (Wang et al. 2007) and SINTAX (Edgar 2016) use simpler and faster word-counting strategies. With these considerations in mind, I designed SINAPS (Simple Non-Bayesian Attribute Prediction Software) to predict traits from a marker gene sequence. From an abstract perspective, SINAPS is essentially the same algorithm as SINTAX. A reference database is provided in which each marker gene sequence is annotated with the trait to be predicted. A random subset of words is extracted from the query sequence and used to find the reference database sequence (*top hit*) with most words in common. This process is repeated 100 times and the most frequently occurring top-hit trait is reported as the predicted trait for the query. The number of iterations in which that trait was the top hit is reported as the bootstrap confidence value.

Measuring accuracy of trait prediction presents similar challenges to assessment of taxonomy prediction. Leave-one-out validation as used by the RDP Classifier unrealistically assumes that a typical query has high identity with the reference database and therefore reports a misleadingly high accuracy (Edgar 2016). A more realistic validation should reflect the fact that most species in a typical sample are known only from their 16S sequences, many of which will have low identities with the closest sequence for a species having an experimentally verified trait (Fig. 1). For this work, I therefore used a variant of two-fold cross-validation in which the reference database is divided into two subsets having a given maximum pair-wise identity (*t*) with each other, modeling a scenario where the closest reference sequence to each query sequence has identity *t*. I used *t* = 97%, 95%, 90% and 85% as representative identities encountered in practice. These identities correspond, very roughly, to species, genus, order and phylum (Yarza et al. 2014). If the query identity is 85%, accurate prediction thus requires that the trait is well conserved within a phylum. As a rough guideline, 85% identity is therefore the low end of a “twilight zone” (Rost 1999) where we should expect trait prediction to become ineffective in practice. For testing, I selected a variety of quite different traits including energy metabolism, Gram positive staining, presence of a flagellum, 16S copy number, and number of V4 primer mismatches.

**Fig. 1.**
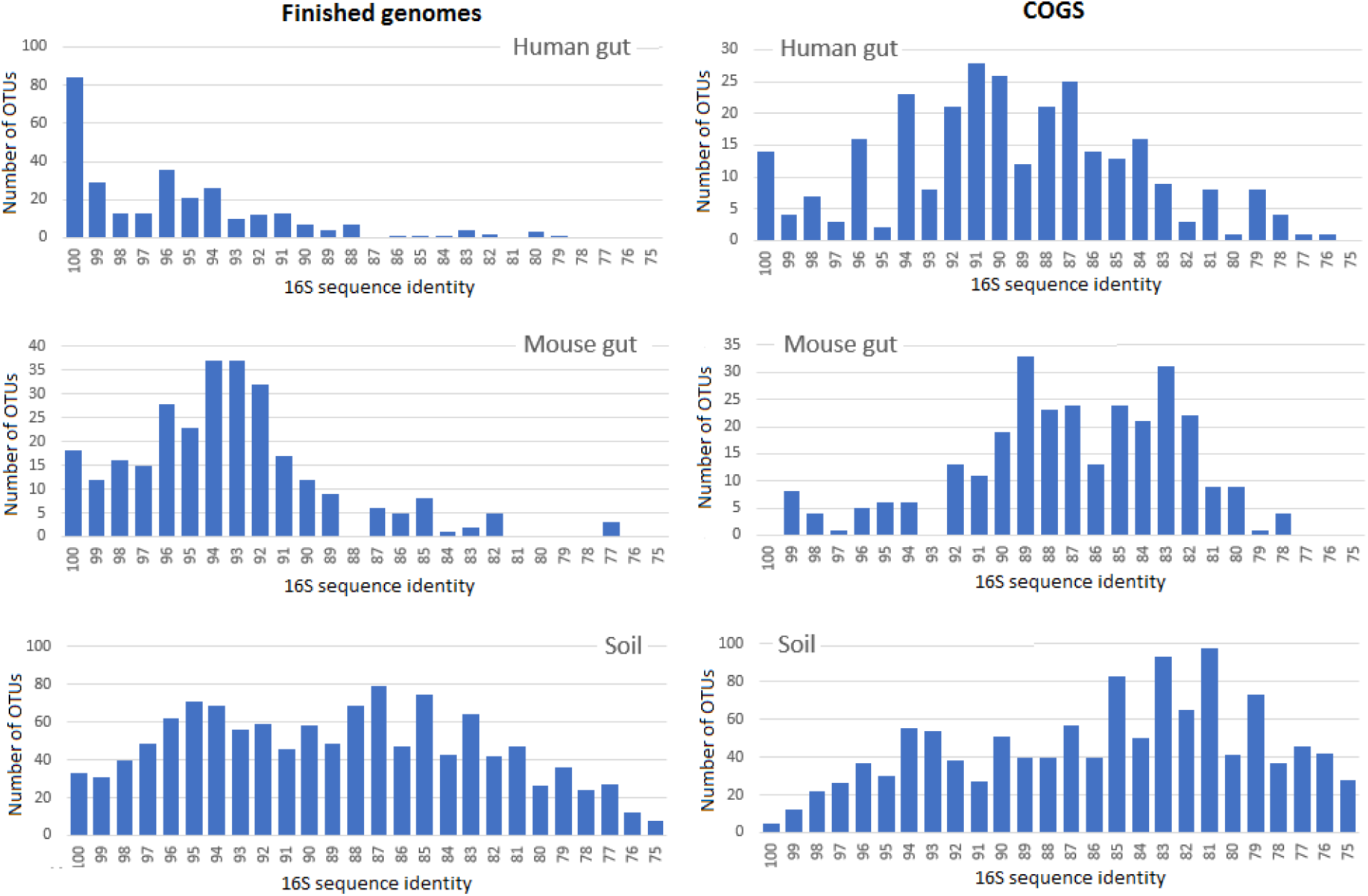
16S sequence identity between OTUs and finished genomes. The histograms show identity distribution for OTUs from human gut, mouse gut and soil samples in a recent study (Kozich et al. 2013) with 16S sequences in the 6,487 currently available finished genomes (left) and the 711 genomes in the Dec. 2014 release of COGS (right).

## Methods

### SINAPS algorithm

Given a query sequence *Q* and reference database *R* annotated with an experimentally determined trait, the SINAPS algorithm proceeds as follows. Let *W(Q)* be the set of *k*-mers in *Q* where *k* = 8 by default. In one iteration, a random sub-sample *Ws*(*Q*) of size *s* is extracted from *W*(*Q*) where *s* = 32 by default. Sub-sampling is performed with replacement. For each reference sequence *r* ∊ *R*, the number of words in common is *U^subset^*(*r*) = |*ws*(*Q*) ∩ *W*(*r*)|. The top hit *T* by *k*-mer similarity is identified as *T* = *argmax*(*r*) *U^subset^*(*r*) and the trait is taken from the annotation of *T*. If there is more than one top hit, *T* is selected at random to avoid the bias that would occur with a systematic rule such as selecting the first in database order, which would give higher confidence to traits found earlier in the database. By default, 100 iterations are performed. The trait that occurs most often is reported as the prediction and its frequency is reported as its bootstrap confidence.

### D16S dataset

To create a reference database of full-length 16S sequences with reliably assigned NCBI taxonomies, I ran the SEARCH_16S algorithm (Edgar 2017) on the 6,487 prokaryotic assemblies in Genbank that were annotated as “Complete genome” as of 15th Jan. 2017, giving a set (D16S) of 22,899 unique sequences.

### 16S copy number

Prokaryotic genomes contain from one to ten or more copies of the 16S gene (Pei et al. 2010), causing bias of an order of magnitude in abundances measured from amplicon reads (Kembel et al. 2012; Edgar 2017). I annotated each D16S sequence with the total number of 16S sequences found by SEARCH_16S in its genome, giving a copy number reference database (D16S-CN).

### V4 primer mismatches

The 16S gene is typically amplified using so-called universal primers that in fact match most, but not all species. Mismatched positions can degrade amplification efficiency by a large factor and are thus another substantial source of amplification bias (Sipos et al. 2007; Edgar 2017). To create a reference database for mismatches to a given primer, it suffices to have full-length sequences, or a partial sequence that covers the segment targeted by the primer. All sequences in a large 16S database such as SILVA or Greengenes could therefore be used to create a reference. For this work, I used the D16S set for simplicity and consistency with other tests as I considered it to be large enough for robust validation. I measured the total number of differences with the currently popular primer pair V4F (GTGCCAGCMGCCGCGGTAA) and V4R (GGACTACHVGGGTWTCTAAT), giving reference set D16S-V4d.

### Energy metabolism

To obtain annotations of energy metabolisms I used the PROTRAITS database (Brbić et al. 2016), selecting the species also found in D16S having integrated metabolism annotations with at least 95% confidence, giving reference set D16S-Energy. PROTRAITS uses the following categories (with corresponding numbers of sequences in D16S-Energy): chemoorganotroph (10,786), heterotroph (1,480), lithotroph (67), methylotroph (24), photoautotroph (5), photosynthetic (36), and phototroph (11).

### Gram-positive staining and presence of flagellum

Gram-positive staining and the presence of a flagellum are binary traits which I again obtained by selecting species present in D16S having integrated PROTRAITS predictions with at least 95% confidence. This gave reference sets D16S-Gram (15,537 sequences) and D16S-Flag (7,705 sequences), respectively.

### Two-fold cross-validation

For each reference set D16S-CN, D16S-V4d, D16S-Energy, D16S-Gram and D16S-Flag I performed two-fold cross-validation as follows. First, a reference set was divided into two subsets *X*_*t*_ and *Y*_*t*_ such that the most similar sequence in the opposite subset has a given identity *t*, discarding sequences as needed to satisfy this constraint. For example, with *t* = 95%, subsets *X*_95_ and *Y*_95_ were constructed so that each sequence in *X*_95_ has a top hit with identity 95% in *Y*_95_, and similarly the top hit in *X*_95_ for every sequence in *Y*_95_ has identity 95%. I created *X*_*t*_, *Y*_*t*_ pairs for *t* = 97%, 95%, 90% and 85% to reflect typical query-reference identities encountered in practice. For each *t*, I measured accuracy for full-length queries by using *X*_*t*_ as a query and *Y*_*t*_ as a reference, and *vice versa*. To assess the use of short tags as query sequences, I extracted the V4 regions from each pair using the primers specified above, giving *X*^V4^_*t*_ and *Y* ^V4^_*t*_. I then used *X*^V4^_*t*_ as a query against the full-length *Y*_*t*_ as a reference and *Y*^V4^_*t*_ as a query against full-length *X*_*t*_.

## Results

Two-fold cross-validation results are summarized in Table 1. Accuracy is defined as the fraction of predictions which are correct. Accuracy is >95% for most traits at most tested identities except for 16S copy number, where the highest measured accuracy is 72.7% (*t* = 97%, ≥90%) bootstrap). This reflects that copy number is not well conserved, especially at lower identities. However, an incorrect prediction that is close to the correct value could still be useful, for example in correcting for amplification bias (Edgar 2017). I therefore also measured the distribution of (predicted copy number) – (true copy number) (Fig. 2). These results show that a large majority of predictions are within +/−2 of the correct value, even at *t* = 85%.

**Fig. 2.**
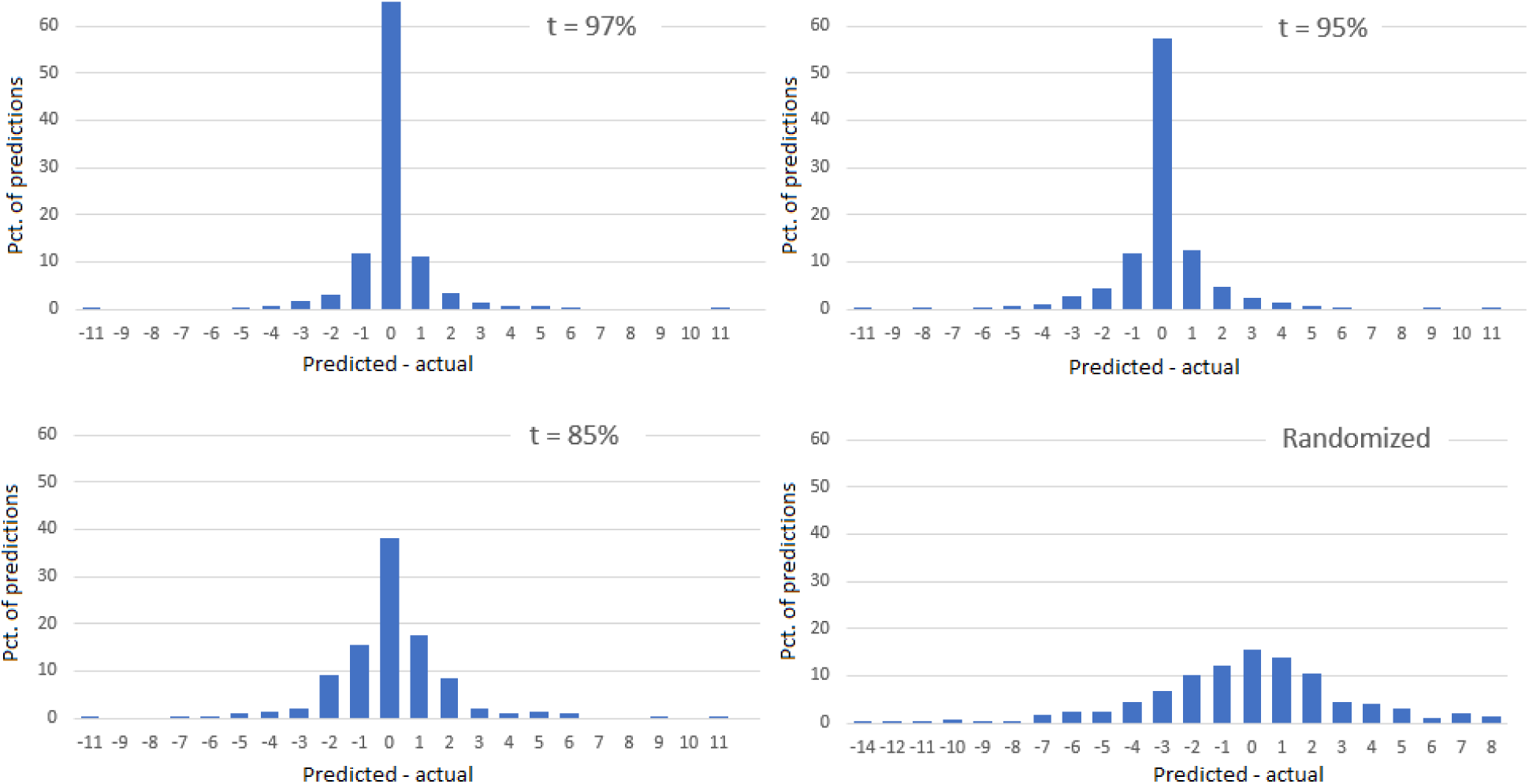
Predicted vs. actual 16S copy number. The histograms show the distribution of (predicted copy number) – (true copy number) for query-reference identity (*t*) 97%, 95% and 85%. The lower-right panel shows the distribution when the predictions are randomized by shuffling, which preserves the frequency of each copy number. This shows that even at 85% identity, predictions are much closer to the correct values than a random guess based on the observed frequencies. Thus, copy number is well-enough conserved at 85% identity (approximately phylum level) to enable useful prediction.

**Table 1.** Two-fold cross-validation results. Accuracy is the fraction of predictions that are correct. *Identity* is the value of *t*, i.e. the top-hit identity between the query and reference database; *Cov (90)* is the fraction of predictions having ≥90% bootstrap; *Acc (90)* is the accuracy of predictions with ≥90% bootstrap; *Acc (all)* is accuracy of all predictions. Accuracies are color coded: dark green >95%, light green >90%, light orange >50%, dark orange <50%. Predictions are >90% accurate for all tested traits except for copy number. See Fig. 2 for further analysis of the copy number predictions.

| **Metabolism** | Full-length |  |  | V4 |  |  |
| --- | --- | --- | --- | --- | --- | --- |
| Identity | Cov (90) | Acc (90) | Acc (all) | Cov (90) | Acc (90) | Acc (all) |
| 85% | 90.0 | 99.4 | 96.2 | 91.4 | 97.1 | 92.3 |
| 90% | 89.9 | 99.1 | 97.9 | 89.7 | 99.3 | 92.1 |
| 95% | 99.0 | 99.3 | 99.1 | 98.6 | 98.8 | 98.7 |
| 97% | 94.9 | 99.2 | 98.4 | 98.1 | 97.8 | 97.7 |

| **Copy nr.** | Full-length |  |  | V4 |  |  |
| --- | --- | --- | --- | --- | --- | --- |
| Identity | Cov (90) | Acc (90) | Acc (all) | Cov (90) | Acc (90) | Acc (all) |
| 85% | 19.7 | 65.2 | 38.0 | 24.7 | 60.6 | 37.0 |
| 90% | 34.5 | 62.2 | 43.3 | 41.2 | 53.6 | 42.6 |
| 95% | 55.3 | 70.7 | 57.4 | 61.6 | 62.1 | 53.4 |
| 97% | 71.9 | 70.7 | 65.0 | 71.8 | 72.7 | 62.9 |

| **V4 pr. mm.** | Full-length |  |  | V4 |  |  |
| --- | --- | --- | --- | --- | --- | --- |
| Identity | Cov (90) | Acc (90) | Acc (all) | Cov (90) | Acc (90) | Acc (all) |
| 85% | 89.9 | 94.2 | 94.2 | 90.7 | 96.7 | 94.7 |
| 90% | 96.1 | 97.6 | 97.0 | 95.8 | 97.8 | 96.9 |
| 95% | 97.5 | 98.9 | 98.4 | 98.7 | 98.7 | 98.5 |
| 97% | 98.4 | 98.4 | 98.8 | 98.8 | 98.9 | 98.6 |

| **Flagellum** | Full-length |  |  | V4 |  |  |
| --- | --- | --- | --- | --- | --- | --- |
| Identity | Cov (90) | Acc (90) | Acc (all) | Cov (90) | Acc (90) | Acc (all) |
| 85% | 76.1 | 71.4 | 73.9 | 73.6 | 75.7 | 73.7 |
| 90% | 72.8 | 90.6 | 70.6 | 68.6 | 94.3 | 75.3 |
| 95% | 97.1 | 99.5 | 98.4 | 97.0 | 98.9 | 98.5 |
| 97% | 97.6 | 99.4 | 98.2 | 98.5 | 97.0 | 96.5 |

**Table 1. Two-fold cross-validation results.**
| **Gram-pos.** | Full-length |  |  | V4 |  |  |
| --- | --- | --- | --- | --- | --- | --- |
| Identity | Cov (90) | Acc (90) | Acc (all) | Cov (90) | Acc (90) | Acc (all) |
| 85% | 94.5 | 99.2 | 98.1 | 91.3 | 99.1 | 97.3 |
| 90% | 98.9 | 99.6 | 99.2 | 97.9 | 99.4 | 99.3 |
| 95% | 99.8 | 99.8 | 99.4 | 99.2 | 99.8 | 99.1 |
| 97% | 99.5 | 99.5 | 99.5 | 99.6 | 99.5 | 99.3 |

## References

Brbić, M. et al., 2016. The landscape of microbial phenotypic traits and associated genes. Nucleic Acids Research, 44(21), pp.10074–10090.

Cho, I. & Blaser, M.J., 2012. The human microbiome: at the interface of health and disease. Nature Reviews Genetics, 13(4), pp.260–270.

DeSantis, T.Z. et al., 2006. Greengenes, a chimera-checked 16S rRNA gene database and workbench compatible with ARB. Applied and environmental microbiology, 72(7), pp.5069–72.

Edgar, R.C., 2016. SINTAX: a simple non-Bayesian taxonomy classifier for 16S and ITS sequences. bioRxiv, p.74161.

Edgar, R.C., 2017. UNBIAS: An attempt to correct abundance bias in 16S sequencing, with limited success. bioRxiv.

Hartmann, M. et al., 2014. Resistance and resilience of the forest soil microbiome to logging-associated compaction. The ISME journal, 8(1), pp.226–44.

Kembel, S.W. et al., 2012. Incorporating 16S Gene Copy Number Information Improves Estimates of Microbial Diversity and Abundance. PLoS Computational Biology, 8(10).

Kozich, J.J. et al., 2013. Development of a dual-index sequencing strategy and curation pipeline for analyzing amplicon sequence data on the miseq illumina sequencing platform. Applied and Environmental Microbiology, 79(17), pp.5112–5120.

Langille, M. et al., 2013. Predictive functional profiling of microbial communities using 16S rRNA marker gene sequences. Nature biotechnology, 31(9), pp.814–21.

McDonald, D. et al., 2012. An improved Greengenes taxonomy with explicit ranks for ecological and evolutionary analyses of bacteria and archaea. ISME J, 6, pp.610–618.

Moran, M.A., 2015. The global ocean microbiome. Science, 347(6219), p.aac8455.

Ogata, H. et al., 1999. KEGG: Kyoto encyclopedia of genes and genomes. Nucleic Acids Research, 27(1), pp.29–34.

Pei, A.Y. et al., 2010. Diversity of 16S rRNA genes within individual prokaryotic genomes. Applied and Environmental Microbiology, 76(12), pp.3886–3897.

Pflughoeft, K.J. & Versalovic, J., 2012. Human microbiome in health and disease. Annual review of pathology, 7(December 2011), pp.99–122.

Pruesse, E. et al., 2007. SILVA: A comprehensive online resource for quality checked and aligned ribosomal RNA sequence data compatible with ARB. Nucleic Acids Research, 35(21), pp.7188–7196.

Rost, B., 1999. Twilight zone of protein sequence alignments. Protein engineering, 12(2), pp.85–94.

Sipos, R. et al., 2007. Effect of primer mismatch, annealing temperature and PCR cycle number on 16S rRNA gene-targetting bacterial community analysis. FEMS Microbiology Ecology, 60(2), pp.341–50.

Tatusov, R.L., Koonin, E. V & Lipman, D.J., 1997. A genomic perspective on protein families. Science, 278(5338), pp.631–637.

Wang, Q. et al., 2007. Naive Bayesian classifier for rapid assignment of rRNA sequences into the new bacterial taxonomy. Applied and environmental microbiology, 73(16), pp.5261–7.

Yarza, P. et al., 2014. Uniting the classification of cultured and uncultured bacteria and archaea using 16S rRNA gene sequences. Nature Reviews. Microbiology, 12(9), pp.635–645.

Yilmaz, P. et al., 2014. The SILVA and “all-species Living Tree Project (LTP)” taxonomic frameworks. Nucleic Acids Research, 42(D1).

